# DeepLabStream: Closing the loop using deep learning-based markerless, real-time posture detection

**DOI:** 10.1101/2019.12.20.884478

**Authors:** Jens F. Schweihoff, Matvey Loshakov, Irina Pavlova, Laura Kück, Laura A. Ewell, Martin K. Schwarz

## Abstract

In general, animal behavior can be described as the neuronal-driven sequence of reoccurring postures through time. Current technologies enable offline pose estimation with high spatio-temporal resolution, however to understand complex behaviors, it is necessary to correlate the behavior with neuronal activity in real-time. Here we present DeepLabStream, a highly versatile, closed-loop solution for freely moving mice that can autonomously conduct behavioral experiments ranging from behavior-based learning tasks to posture-dependent optogenetic stimulation. DeepLabStream has a temporal resolution in the millisecond range, can operate with multiple devices and can be easily tailored to a wide range of species and experimental designs. We employ DeepLabStream to autonomously run a second-order olfactory conditioning task for freely moving mice and to deliver optogenetic stimuli based on mouse head-direction.

## Introduction

Recent developments in the field of behavioral research have made offline pose estimation of several species possible using robust deeplearning-based markerless tracking ^1,2^. DeepLabCut (DLC), for example, uses trained deep neuronal networks to track the position of user defined body parts and provides offline motion tracking of freely moving animals. Additionally, sophisticated computational approaches have allowed for disentangling the complex behavioral expressions of animals into patterns of reoccurring modules ^3–5^. *In vivo* single unit recording ^6^, along with recent advances in *in vivo* voltage imaging ^7^ and miniaturized calcium imaging techniques ^8–10^, facilitate realtime measurements of neuronal activity in freely moving animals. Together, these techniques provide a platform for correlating recorded neuronal activity and behavior. Traditionally, to test the correlations, either neuronal activity or behavior would be manipulated experimentally. Classic manipulations of neuronal activity such as lesions, transgenic alterations, and pharmacological injections result in long-lasting, and sometimes chronic changes in the tested animals, which can make it difficult to interpret behavioral effects. In recent years, there has been a shift towards techniques that allow for fast, short-lived manipulation of neuronal activity. Optogenetic manipulation, for example, offers high temporal precision, enabling the manipulation of experience during tasks that test mechanisms of learning and memory ^11–13^, perception^14,15^ and motor control ^16,17^. Such techniques offer a temporal resolution precise enough that the neuronal manipulation can match the timescale of either behavioral expression or neuronal computation. For best effectiveness, however, behavior must be identified in real-time, allowing for instantaneous feedback, i.e. “closed-loop” manipulation based on this behavioral expression ^18,19^. Currently, such experimental systems often rely on specialized, on-purpose setups, including intricate beam brake designs, treadmills, and virtual reality setups to approximate the movement of the investigated animal in a given environment and then react accordingly ^20–26^.

We here introduce DeepLabStream (DLStream), a multi-purpose software solution that enables markerless, real-time tracking and neuronal manipulation of freely moving animals during ongoing experiments. Its core capability is the orchestration of closed-loop experimental protocols that streamline posture-dependent feedback to several input-, as well as output-devices. We modified state-of-the-art pose estimation based on DLC ^1^ to be able to track the postures of mice with body part precision in real-time. We demonstrate the software’s capabilities in a classic, multilayered, freely moving conditioning task, as well as in a head direction-dependent optogenetic stimulation experiment using an activity-dependent, light-induced labeling system ^18^. Finally, we show the versatility of DLStream to adapt to different experimental conditions and hardware configurations.

## Results

DLStream enables closed-loop applications directly dependent on behavior expression. Our solution is fully autonomous and requires no additional tracking-, trigger- or timing-devices. All experiments can be conducted without any restriction to the animal’s movement and each experimental session is run fully autonomously after the initial setup. Initially, we trained DLC-based pose estimation networks offline for each experimental environment and then integrated them into DLStream (see Materials and Methods). Briefly, frames were taken from a video camera stream and analyzed using an integrated deep neuronal network, trained using the DLC framework. Next, the resulting pose estimation was converted into postures and transferred to an additional process that supervises the ongoing experiment and outputs feedback to connected devices (See Figure 1). As a result, experiments run by DLStream comprise a sequence of modules (see Figure 2 C) depending on the underlying experimental protocol. Basic modules, such as timers and stimulations, are posture-independent and control fundamental aspects of the experiment. Timers keep track of the time passing during frame-by-frame analysis and act as a gate for posture-based triggers and stimulations (e.g. interstimulus time). Stimulations specify which devices are triggered and how each device is controlled once it was triggered (e.g. reward delivery). Posture-based triggers are sets of defined postures (e.g. position, head-direction, etc.) that initialize a predefined cascade (stimulation) once detected within an experiment (see Figure 2 for examples). As an experiment is conducted, DLStream records and subsequently exports all relevant information, including posture tracking, experimental status and response latency in a tablebased file. During any experiment, the posture tracking is visualized on a live video stream directly enabling the monitoring of the conducted experiment and tracking quality. Additionally, the raw video camera stream is timestamped and recorded, allowing high-framerate recording, with lower-framerate closed-loop posture detection to save processing power (See Figure 1).

**Figure 1.**
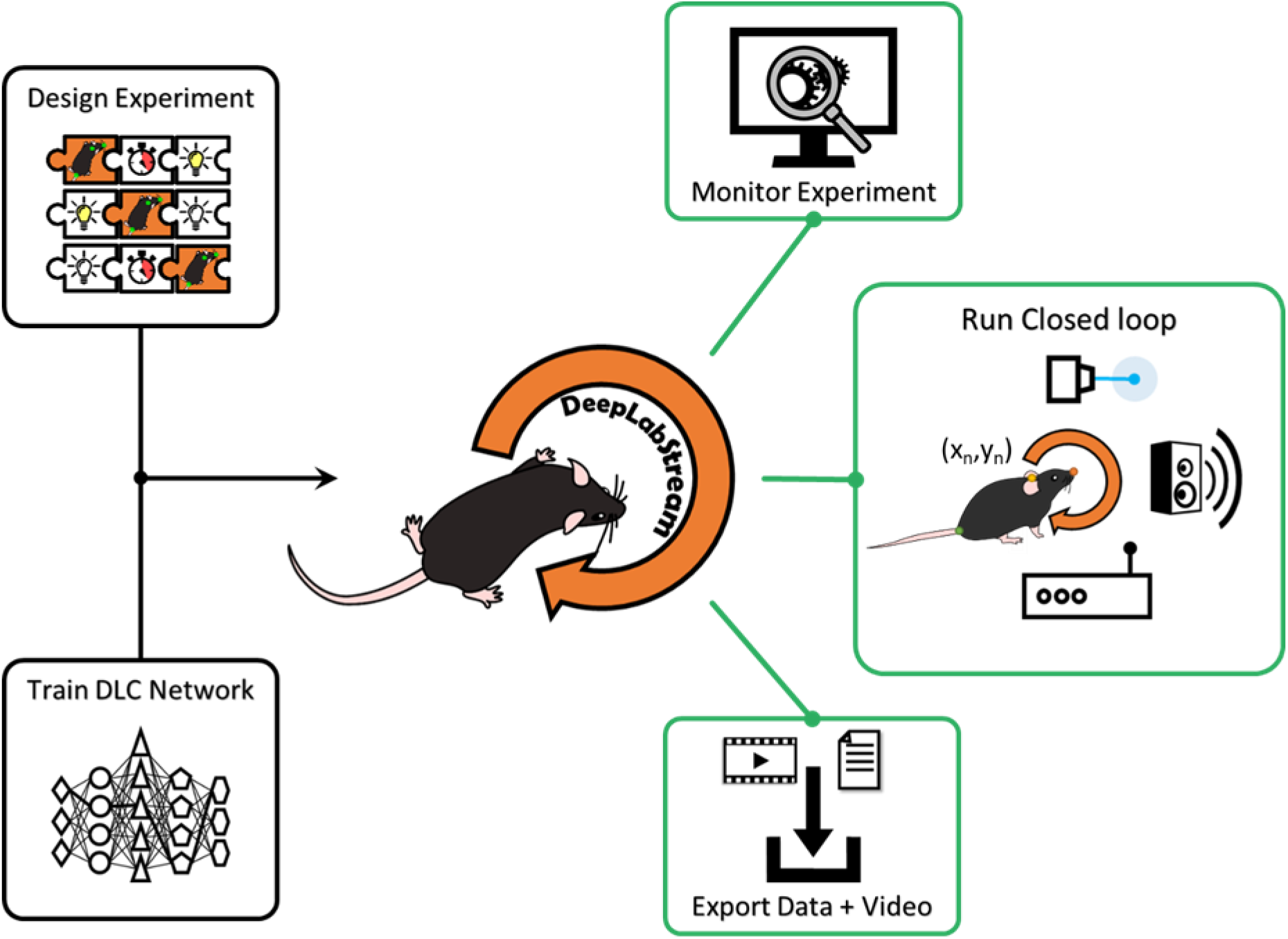
A visual representation of DLStream. Visual representation of workflow in DLStream. Initially, an experimental protocol is designed using a sequence of modules (puzzle pieces) and a trained DLC network is integrated into DLStream. Afterwards, DLStream provides three different outputs for every experiment. 1. Experiments can be monitored on a live stream. 2. The experimental protocol is run based on posture detection 3. Recorded video and experimental data are exported after the experiment are done.

**Figure 2.**
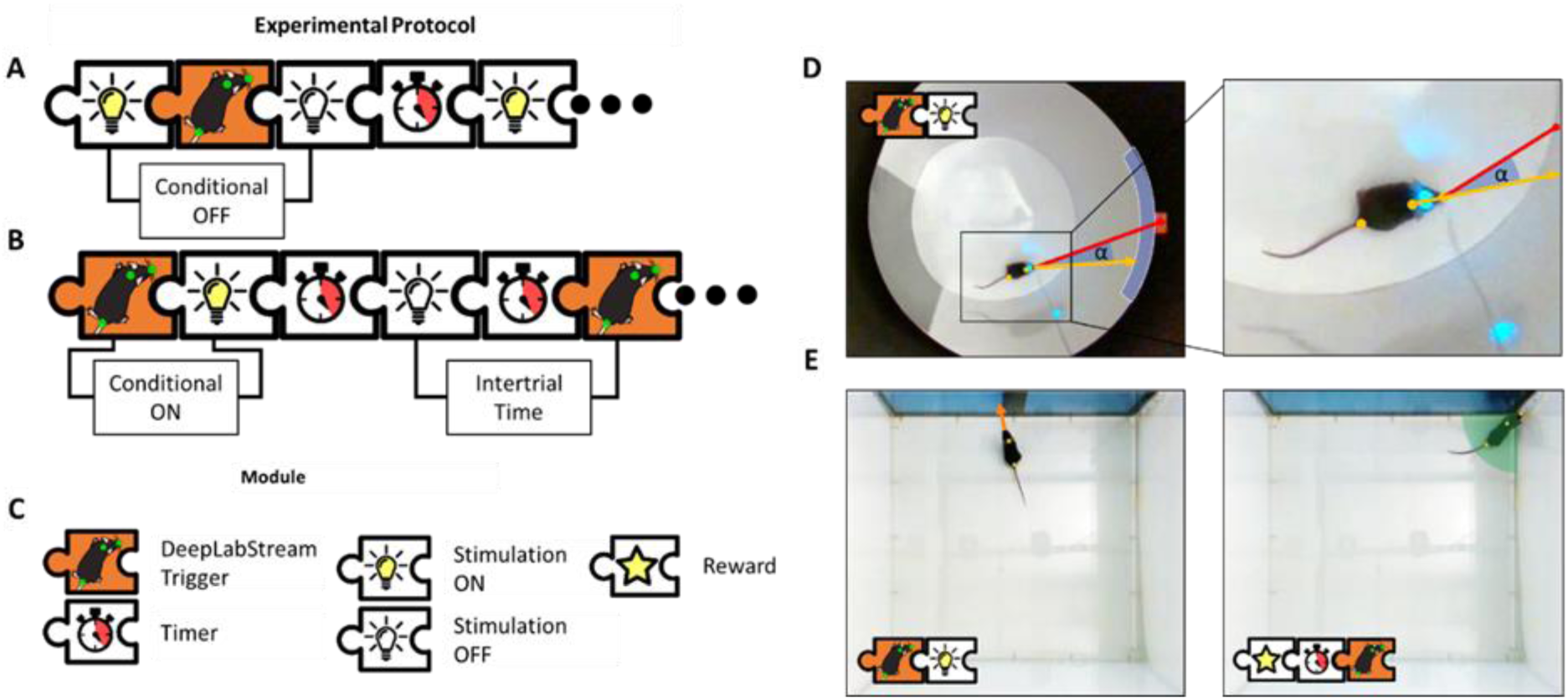
Experimental Design using DLStream. (**A, B**) Schematic design of an experimental protocol with a posture-based trigger. Manipulation can be turned OFF (**A**) and ON (**B**) based on the mouse’s behavior. The combination of several modules allows building a sophisticated experimental protocol. (**C**) Description of available modules in (**A**) and (**B**). (**D**) Application of the above-described design in an optogenetic experiment. The stimulation is triggered dependent on head direction angle (orange arrow, α) to a reference point (red line) within window (blue arc). (**E**) Application of the above-described design in a classical conditioning task. The mouse is shown an image when looking at the screen (left) and the reward is removed if it does not move into the reward location within a timeframe (right, green zone). The mouse’s posture is shown with orange dots.

### Classical second-order conditioning using DLStream

To comprehensively test DLStream we first designed a fully automated classical second-order conditioning task (Figure. 3, A-E). Using DLStream, mice were trained to associate two unknown odors (rose, vanillin) with two visual stimuli, which were initially associated with either a reward or an aversive tone (Figure. 3, A). We subsequently tested the conditioned mice in an odor preference task.

**Figure 3.**
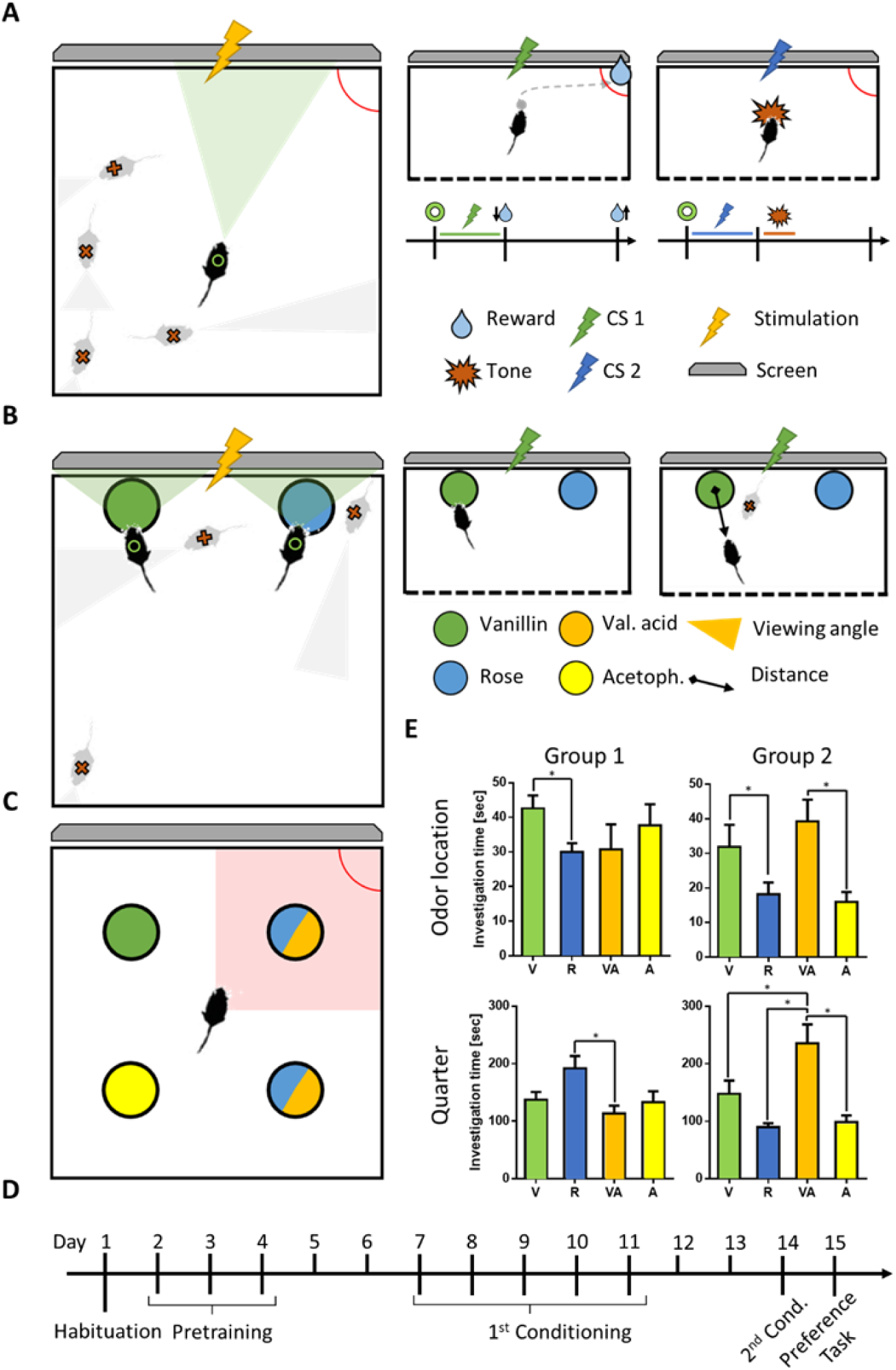
Closed-loop conditioning task. **(A)** Conditioning. When a trial is triggered by the mouse facing the screen (left, green triangle and ring), the mouse is shown a visual stimulus (yellow lightning bolt). Mice not facing the screen do not receive the stimulus (red x, yellow triangle). In the positive trial (middle; green lightning bolt, green line), a reward is delivered (blue drop, arrow down) and withdrawn (blue drop, arrow up) if not collected within a preset time period. In the negative trial (right; blue lightning bolt, blue line) only a loud tone (red polygon) is delivered. **(B)** 2nd Order Conditioning. Upon exploration of either odor location (colored black circle) the mouse is shown one of the previously conditioned visual stimuli on the screen (left, yellow lightning bolt). Conditioning was conducted in two stages (top middle and top right). The first stage (top middle) consisted of direct contact with the odor location, while the second (top right) was dependent on the proximity of the mouse to one of the locations (black arrow) and the mouse facing towards it. **(C)** Odor Preference Task. The mouse was set in an open field arena with one odor in each of the quarters (colored circles). The multicolored circles represent the differences in odor location between the two groups. The red square shows the previous reward location and quarter. (**D**) Schematic of experimental workflow: The timeline is represented as days (numbers on top) and experimental stages (below) with their corresponding order and length (brackets for multiple days). **E**: Investigation time during odor preference task measured in two regions of interest (ROI). Odor location: ROIs encircling the odor location (**A, B**). Quarter: Rectangular ROIs dividing the arena into quarters (**C, D**). Group 1 had Rose odor placed in the reward collection quarter, while Group 2 had valeric acid placed in it. p < 0.05 (*) Odor location: paired t-test; Quarter: 1-way ANOVA. Error bars show SEM. n = 5 per group. V = Vanillin (S+), R = Rose (S-), VA = Valeric acid, A = Acetophenone

In the first conditioning stage, DLStream triggered trials when a mouse was facing the screen. For this, a trigger module was designed that utilizes the general head direction of mice, activating stimulation modules only, when mice were looking towards the screen in a 180° window. The mice were conditioned to associate two unknown visual stimuli (a high contrast black and white image) with a reward or an aversive tone (Figure. 3, A) using combinations of predefined stimulation modules. In the positive trial, DLStream delivered a liquid reward by triggering the corresponding stimulation module in a fixed reward location and withdrew it if it was not collected within a preset time period monitored with a timer module. In the negative trial, DLStream delivered only the aversive tone (Figure. 3, A). All mice were trained for 4-6 days depending on individual performance with an average of 213 ± 30 trials to reach the success criterion (85 % reward collection within one session, n = 10). We limited the number of sessions to 1 h or 40 trials per day within our experimental protocol. Note that no mouse needed more than 45 min to complete a session. During the subsequent second-order conditioning, the mice were presented with two novel odors (rose and vanillin), placed in a petri dish in front of the screen (Figure. 3, B). Visual stimuli were previously paired with an odor and pairing was kept throughout all experiments. Upon exploration of one of the two presented odors, DLStream showed the mice the paired, previously conditioned visual stimulus (Figure. 3, B). The session was completed when DLStream detected that the mice had explored both odors at least 10 times, or after 10 minutes had passed. Second-order conditioning was then conducted in two stages. The first stage consisted of the mouse being in direct contact with the odor location (petri dish), while the second was dependent on the proximity of the mouse to one of the locations and its head direction (Figure. 3, B). For this, trigger modules designed to detect proximity and heading direction of mice were used. Each stage was repeated twice with exchanged odor locations.

We then tested for successful second-order conditioning by conducting an offline odor preference task (Figure. 3, C). Mice were placed in an open field arena with one odor in each of the quarters. In addition to the two conditioned odors, two novel odors (acetophenon and valeric acid) were presented. Mice were given 10 min to explore and total investigation time was measured. The investigation time was measured by taking the circular odor location as well as the quarters of the arena into account. Mice showed a clear preference towards the positive conditioned (S+) odor compared to the negative conditioned (S-) odor, spending significantly more time at the S+ odor location than in the Sodor location (Figure. 3, E). Interestingly, we noticed differences in investigation time when considering the full quarter of the arena rather than the odor location itself. Therefore, we tested whether odor placement within the arena had a significant influence on preference. We separated the mice into two groups. One group had a neutral odor placed in the reward quarter (top right), while the other group had the Sodor at that position. A novel odor placed in the quarter where the reward location was in the previous conditioning task, had significantly higher investigation time than all other odors (one-way ANOVA Group 2; A/VA p = 0.003, R/VA p = 0.0001, V/Va p = 0.0248 in quarter ROI), while there was no significant difference between both conditioned odors when the Sodor was placed in the reward quarter (Figure. 3, E) (one-way ANOVA Group 1; R/VA p = 0.0146, rest ns. in quarter ROI).

### Optogenetic, head-direction-dependent labeling of neurons using DLC

As a second example of the wide applicability of DLStream, we optogenetically stimulated mice injected with the neuronal activity indicator Cal-Light, within precisely defined head direction angles (Figure. 4 A, B; data not shown). Mice were placed in a circular white arena with a single black cue at one side and allowed to investigate the arena in 30-minute sessions per day for at least 5 consecutive days. During each session, mice were stimulated via a chronically implanted optical fiber with blue light (488 nm) dependent on the head direction angle. Stimulation was limited to the designated head direction window (60° to reference point, Figure 4 D, E), while mice freely moved their heads in all directions (Figure 4 F). Mice explored the complete arena during the task. Note that the resulting light stimulation was independent of the animal’s position in the arena (Figure 4 G). As a control we show that the head direction-dependent stimulation cannot be reproduced by stimulation at random time points during the session (Figure 4 H) and is significantly different from the distribution of average head direction angles by random sampling (n = 10000, p < 0.01). Each stimulation lasted 1-5 sec depending on the time spent in the pre-defined “correct” head direction window (60°) with a minimum inter-stimulus time of 15 seconds. Timing was controlled by designated timer modules controlling onset and offset of light stimulation once the stimulation module was triggered. In the case of the interstimulus timer, the module blocked the link between trigger module and stimulation module when activated, disabling posture-dependent stimulation for its designated duration.

**Figure 4.**
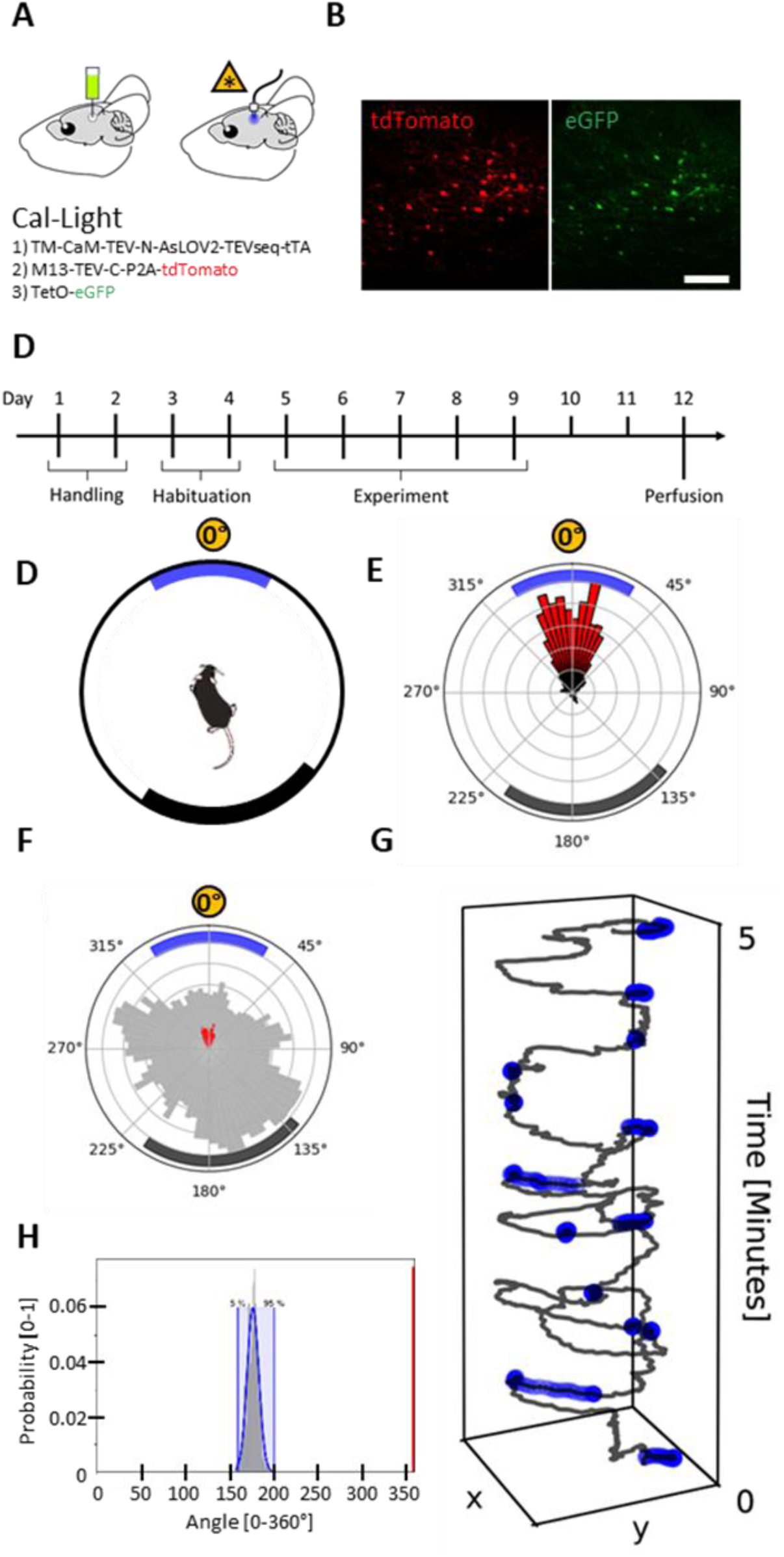
Optogenetic labeling of head direction-dependent activity in neurons. **(A)** Stereotactic delivery of Cal-Light components into the forebrain and fiber ferrule placement (left) as well as stimulation above the region of interest (right). (**B**) Example of viral targeting with tdTomato expression (left, red) and eGFP expression (right, green). The bar represents 100 µm. (**C**) Schematic of experimental workflow: The timeline is represented as days (numbers on top) and experimental stages (below) with their corresponding order and length (brackets for multiple days). (**D**) Scheme of the circular arena with the cue (thick black arc) and the stimulation window (thick blue arc) around the reference point (orange circle). (**E**) Representative example radial histogram of all head directions of a mouse during stimulation (red) within one session. Normalized to the maximum value. Rings represent quantiles in 20 % steps. (**F**) Radial histogram of all head directions of the same mouse and session as in (**D**) during the whole session (grey) and during stimulation (red). Normalized to the maximum value of the whole session. Rings represent quantiles in 20 % steps. (**G**) Representative example of the mouse’s position (grey) over time during the first 5 minutes of the session in (**E, F)**. The stimulation events are shown in blue. (**H**) Distribution of average head direction in random sampling (n = 10000) from the session shown in (**F**) with 5% and 95% percentile in blue. The red line denotes the actual mean head direction during stimulation in the session (**E**).

### Computational Performance of DLStream

A reality of any closed-loop system is that there are temporal delays between realtime detection of particular postures and stimulus output. To address this challenge, we first rigorously defined the variance of behavioral parameters we are measuring. To estimate the spatiotemporal resolution of postures that can be detected using our integrated network configuration, we compared the displacement of individual body parts between frames. DLC network performance was 2.28 pixels (ca. 3 mm in our setup) using the provided DLC evaluation tools (see Materials and Methods). Within our experiments (n = 21) the average Euclidean distance between two consecutive frames was 2 ±3 mm measured at the neck point and 2 ± 4 mm at the tail root, while nose point displacement was 4 ±9 mm (Figure 5 B-F). Body part estimation resulted in an average head direction variance of 11.8 ± 47.5° (n = 21) between consecutive frames. Note that the variance is a product of performance errors and the inhomogeneous movement of the animal during experiments. Depending on the mixture of episodes of fast movements and slow movements during sessions, the deviation might change.

**Figure 5.**
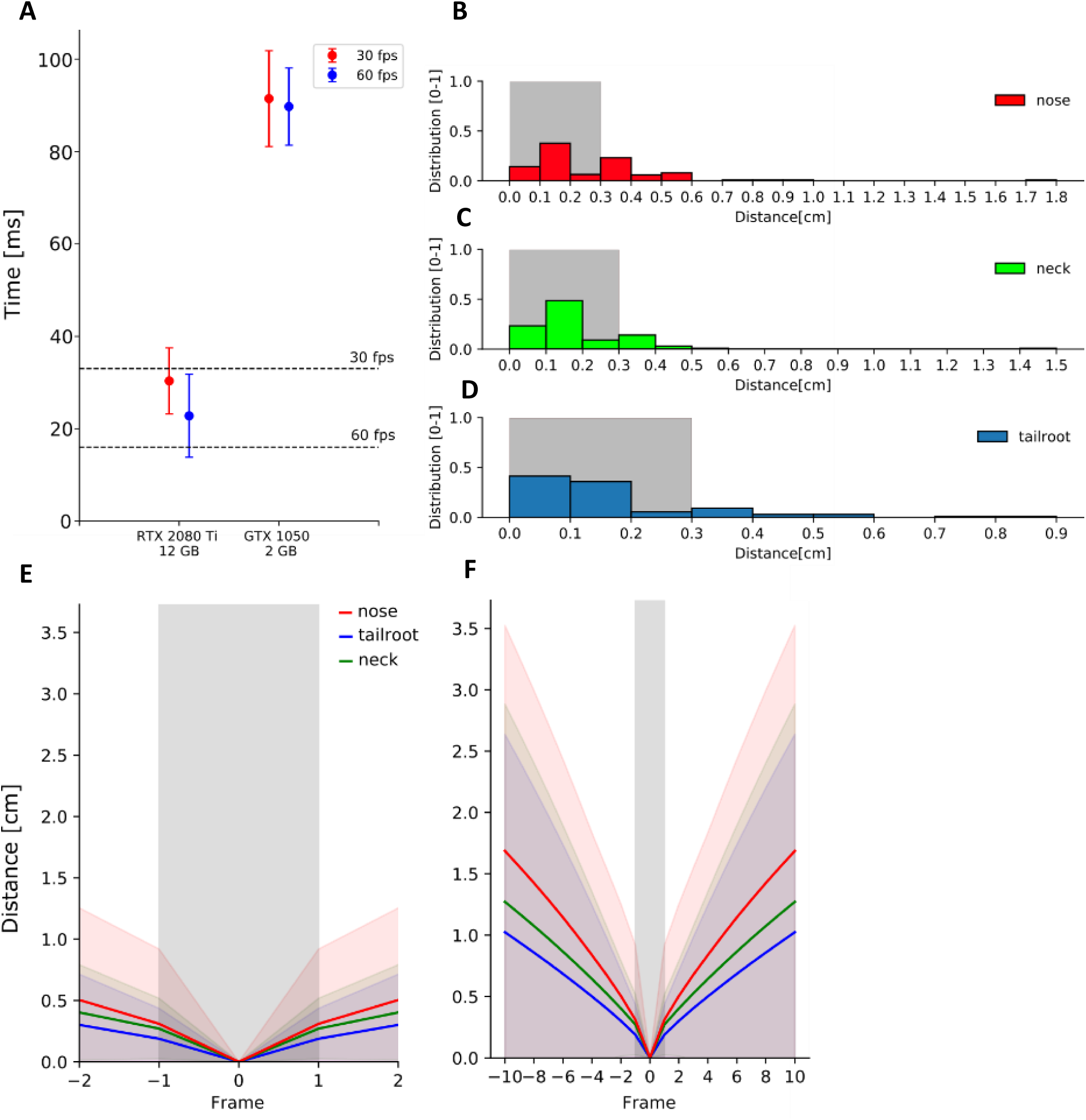
Average Real-time Performance. (**A**) Average performance (n= 10000) of real-time DLStream analysis in two different configurations with 30 fps and 60 fps with 848×480 resolution. The dotted lines represent the anticipated average framerate 30 and 60 with 33.33 ms and 16.66 ms respectively. (**B-D**) Representative distribution of the Euclidean distances between two sequential frames of the same body part compared to the estimated network error (grey window). (**E, F**) Representative average Euclidean distance between sequential frames of the same body part. The x-axis shows the spacing of frames in a sequence that were used to calculate the change in distance (e.g. Distance between frames that were 2 frames apart are depicted at -2 and +2). The colored areas show the standard deviation. The average Euclidean distance the chosen framerate is able to resolve is highlighted in grey.

We tested different hardware configurations to investigate performance levels and response time. Average performance was measured during 10000 frames in two different configurations with two different camera settings (30 fps and 60 fps with 848×480 px resolution) using the same camera (see Figure 5 A). Note that we tested the response time only on a software level, as the total latency depends on individual setups and intrinsic parameters of connected components. With the standard 30 fps camera setting, the advanced configuration (Intel Core i7-9700K @ 3.60GHz, 64 GB DDR4 RAM and NVidia GeForce RTX 2080 Ti(12GB) GPU) achieved reliable 30 fps (33.33 ms per frame) real-time tracking with 30 ± 7 ms, while the other system (Intel Core i7-7700K CPU @ 4.20GHz, 32GB DDR4 RAM and NVidia GeForce GTX 1050 (2GB) GPU) only reached an average analysis time of 91 ± 10 ms. Using a higher framerate input from the camera (60 fps; 16.66 ms per frame), the overall performance did not change significantly (24 ± 9 ms and 90 ± 9 ms respectively). During optogenetic experimental sessions (n = 87), DLStream reached an average performance time of 32.75 ± 0.36 ms per frame, matching the average camera framerate of 30 Hz (33.33 ms). This includes posture detection and computation of resulting experimental protocols until the output is generated.

## Discussion

### DLStream experimental design

There has been a recent revolution in markerless pose estimation using deep neuronal networks. However, a limitation of these systems is their intrinsic design delays analysis until after the end of the experiment owing to their heavy computation. Here we take advantage of the power of DLC’s offline body part tracking and redeploy it as a real-time, closed-loop solution.

As observers, experimenters often record and interpret an animal’s behavior by taking its movement as an approximation of the underlying intention or state of mind. Building on this generalization, behavior can be defined, categorized and even sequenced by examining estimations of the animal’s movement ^4,5,27,28^. Classified periods of behavior, so called behavior modules, are commonly used for offline quantification (e.g. phenotyping). But behavior modules are also very promising in closed-loop approaches to react specifically to a complex behavior. Such an analysis yields the prospect of predicting behavior, for example by matching initial elements of a uniquely-arranged behavioral sequence. With DLStream, a combination of triggers based on the animal’s posture or posture sequence is now possible. Example triggers include center-of-mass position, direction, and speed of an animal, although multiple individual tracking points can also be utilized, such as the position and trajectory of multiple body parts. This allows the design of advanced triggers that include head direction, kinematic parameters, and even specific behavior modules (e.g. rearing, grooming or sniffing). Out of the box, DLStream supports triggers based on single-frame postural information, although posture sequences or complex behavior modules are also possible once behavior based on collected posture data has been classified, modeled, and integrated as trigger modules into DLStream.

Two of the most significant limitations in all real-time applications are the latency of the system to react to a given input and the rate in which meaningful data is obtained. While the latency is dependent on the computational complexity, the rate dependent on several factors, and hardware constraints in particular. A researcher might only need the broadest movements or behavioral states to understand an animal’s basic behavior, or fast, accurate posture sequences to classify behavioral modules on a sub-second scale ^5,27^. Considering that animals behave in a highly complex manner, a freely moving approach is favorable since restricting movement likely reduces the read-out of the observable behavioral spectrum.

DLStream is designed as a universal solution for freely moving applications and can, therefore, be used to investigate a wide range of organisms. DLC networks already have the innate capability to track a variety of animals across different species ^29^ which can be directly translated to experiments within DLStream. Additionally, as DLStream is a universal solution, its architecture was designed for short to mid-length experiments (minutes to hours). There are no built-in limitations to conduct long-lasting experiments (days to weeks), but DLStream currently lacks the capability to automatically process the large amounts of raw video data or other utilities that become necessary when recording for longer periods of time. One possible solution would be to remove the raw video output and only save the experimental data that includes posture information, which would considerably lighten the necessary data storage space.

With regards to latency, the current achievable, closed-loop timescale enables the tracking and manipulation of a wide range of activities a rodent might perform during a task (see Figure 5). Very fast movements, however, like whisker movement ^30,31^ and pupil contraction ^32,33^ might not be fully detected using a 30 Hz configuration. Most freely moving applications usually lack the resolution to visualize whiskers and pupils while maintaining an overview of the animal’s movement in a large arena. Note that offline analysis of raw, higher framerate videos can still be recorded if desired. DLStream is able to take frames from a higher framerate stream but still maintain a lower, loss-less closed-loop processing rate (see Figure 5). On a side note, developments in alternative, non-video-based, specialized tracking (e.g. eye-tracking; ^34^) might lead to a solution for researchers interested in capturing truly holistic behavioral data.

### DLStream performance in a Classical Conditioning task

Using posture-dependent conditioning, mice were able to successfully learn an association between a visual stimulus and a reward, thus demonstrating DLStream’s capabilities with respect to the full automatization of classical learning tasks. Second-order conditioning resulted in a clear odor preference when compared to two novel odors. Importantly, mice did not need any previous training apart from initial habituation to the reward delivery system to perform this task. Our results suggest that mice developed a place preference during the first conditioning that was translated into the odor preference task. Mice had a strong preference for the previous reward quarter, seemingly independent of novelty or association of the placed odor. Additional experiments are necessary to further investigate the underlying mechanisms. Never the less, mice showed a clear odor preference towards the positive conditioned odor. Importantly, possible applications are not limited to classical conditioning tasks. Many behavioral tasks, an operant conditioning task for example, could also be accomplished by setting a specific posture or sequence of postures as a trigger to reward, punish or manipulate freely-behaving animals during an experimental session.

### Optogenetic, posture-dependent stimulation of neurons using DLStream

To dissect and better understand the neuronal correlates of complex behaviors an understanding of the actively participating neuronal assembles is desirable. Techniques that can bridge connectomics, electrophysiology, and ethology hold the potential to reveal how computations are realized in the brain and subsequently implemented to form behavioral outcomes. For instance, by utilizing activity-dependent labeling systems such as Cal-Light ^18^, Flare ^19^ or Campari ^35^ it is already possible to visualize active neurons during episodes of behaviors of interest. However, the identification of episodes and following activation of a specific trigger is mostly restricted by a lack of dynamic closed-loop systems. With DLStream, we show that the realtime detection of specific behaviors in freely moving mice can be combined with activity-dependent labeling systems to investigate the neuronal correlates of behavior. We here stimulated neurons in the forebrain of mice depending on their head direction, thereby only targeting ensembles of active neurons ^18^. Stimulation was restricted to a defined head direction angle, while mice moved freely in all directions through the arena. Demonstrating that DLStream can potentially label specific ensembles of active neurons during relevant behavioral expressions. Of course, direct optogenetic activation and inhibition ^36–39^ of neuronal population based on posture-detection is also possible with DLStream. Using a solution like DLStream the range of detectable behaviors increases substantially and applications for activity- and posture-dependent labeling and subsequent manipulation of different freely moving species are wide ranging.

### Performance improvements and Compatibility

Comparative tests between our available computer configurations suggest that the GPU power is responsible for major performance gains in real-time tracking utilizing DLStream. CPU power is also important since several parallel processes need to be maintained during complex experimental protocols and processing of pose estimation. DLStream is able to analyze new frames as soon as the current frame is fully processed, therefore a higher framerate does not slow down DLStream but rather enables it to work at the upper-speed limit (see Figure 5). At this stage, the full utilization of higher framerates will heavily depend on the hardware configuration. Although, further testing is necessary to establish whether a more advanced computer system will provide support for framerates higher than 60 Hz. Apart from the framerate, image resolution has been a major limitation for offline pose estimation using the DLC trained networks. Considering that DLC is able to scale with available GPU power, we approximate that a higher resolution can be made available in DLStream using further advanced configurations without any significant increase in latency.

DLStream is compatible with old and new versions of DLC. Although originally developed for DLC 1.11 (Nature Neuroscience Version, ^1^), we have successfully tested the newest DLC version (DLC 2.x, ^1^) without encountering any problems. Networks trained on either version can be fully integrated into DLStream and used as needed. Additionally, DLStream is theoretically able to support positional information from other pose estimation networks ^2,40^. Recent work from Graving et al. ^40^ suggests that using a different network architecture similar result with faster processing speed can be achieved in offline analysis. Whether this can be directly translated into closed-loop performance needs to be tested, especially considering the quality of tracking.

Notably, DLStream could also be easily upgraded to use 3D posture detection. To achieve this, two reasonable approaches exist that allow 3D tracking of animals based on video analysis. A DLC native approach would be the use of multiple camera angles to triangulate the animal’s position (see ^29^ for further information). An alternative approach would be to use of depth cameras to estimate the distance of an animal to the camera and thereby generate a 3D representation.

## Conclusion

DLStream is a highly versatile, closed-loop software solution for freely moving animals. While we show its applicability in posture-dependent learning tasks and optogenetic stimulation using mice, there are no limitations to the applicability of DLStream on different organisms and other experimental paradigms.

## Code availability

DLStream will be made available to the scientific community upon peer-review.

## Acknowledgements

We would like to thank Jon Ewell for language editing and feedback on the manuscript. We also want to thank Liubov Sokhranyaeva for assistance establishing the Cal-Light system. Work was supported by the DFG, SFB 1089, SPP 2041 to MKS and VW Stiftung to LAE.

## Methods

### Animals

C57BL/6 mice were purchased from Charles River (Sulzfeld, Germany) and maintained on a 12-h light/12-h dark cycle with food and water always available. All the experiments were carried out in accordance with the German animal protection law (TierSCHG), FELASA and were approved by the animal welfare committee of the University of Bonn.

### Surgical procedure

Viral injections were performed under aseptic conditions in 2 months old C57BL/6 mice. Mice were initially anesthetized with an oxygen/isoflurane mixture (2%–2.5% in 95% O2), fixed on the stereotactic frame, and kept under a constant stream of isoflurane (1.5%–2% in 95% O2) to maintain anesthesia. Analgesia (0.05 mg/kg of buprenorphine; Buprenovet, Bayer, Germany) was administered intraperitoneal prior to the surgery, and Xylocaine (AstraZeneca, Germany) was used for local anesthesia. Stereotactic injections and implantations of light fiber ferrules were performed using a stereotactic frame (WPI Benchmark/Kopf) and a microprocessor-controlled minipump (World Precision Instruments, Sarasota, Florida). The viral solution was injected bilaterally into the forebrain. To reduce swelling animals were given Dexamethason (0.2 mg/kg). For implantation, the skin on the top of the scalp was removed and the skull cleared of soft tissue. Light fiber ferrules were implanted and fixed using a socket of dental cement. Lose skin around the socket was fixed to the socket using tissue glue (3M Vetbond). Directly after the surgery animal were administered 1 ml 5% Glucosteril solution. To prevent the wound pain, analgesia was administered on the three following days. Animals were left to rest for at least one week before starting handling. Experiments were conducted three weeks after surgery.

### Perfusion

The mice were anesthetized with a mixture of Xylazine (10 mg/kg; Bayer Vital, Germany) and ketamine (100 mg/kg; Bela-pharm GmbH & Co. KG, Germany). Using a peristaltic pump (Laborschlauchpumpe PLP33, Mercateo, Germany), the mice were transcardially perfused with 1× PBS followed by 4% paraformaldehyde (PFA) in PBS. Brains were removed from the skull and post-fixed in 4% PFA overnight (ON) at +4°C. After fixation, the brains were moved into PBS containing 0.01% sodium azide and stored at +4°C until sectioning. Fixed brains were sectioned coronally (70 or 100 μm) using a vibratome (Leica VT1000 S) and stored in PBS containing 0.01% sodium azide at +4°C.

### Conditioning Task

Animals were placed in an open field arena (70×70 cm). Each session lasted 1h or a maximum number of 40 trials. A session consisted of a random sequence of trials. Additionally, if an animal successfully finished 20 positive trials, the session was ended. A trial was initiated when the animal was facing the screen. Each trial lasted 20 seconds with an inter-trial interval of 30 seconds. At the beginning of each trial, a visual stimulus was shown on the screen for 10 seconds. In the positive trial, a reward was delivered at the end of the visual stimulus and withdrawn if not collected within 7 seconds. In the negative trial, a loud tone (100 dB) was delivered and no reward was given. After at least 5 sessions, animals that learned the association successfully (>85% success rate in the positive trial) were transferred to the next stage.

The visual stimulus was a high-contrast, black and white image of an X or + spanning the whole screen. The screen was as the same size as the arena wall it was placed at.

### Second-order conditioning task

Animals were placed in the open field arena. Two petri dishes filled with fresh bedding were placed on the wall facing the screen. Two odorants (10 µl on filter paper) were placed in one of the petri dishes each. A pair of an odorant and visual stimulus (negative or positive) was chosen and kept throughout the experiments. Upon exploration of an odor location, the animal was shown the corresponding visual stimulus. The session was completed after the animal explored both odors for at least 10 individual times or after 10 minutes. Conditioning was conducted in two stages. Both stages were repeated with switched odor positions, resulting in a total of 4 repetitions per animal. The first stage consisted of direct contact with the odor location, while the second was dependent only on the proximity of the animal to a location and the animal facing towards it.

### Preference task

The animal was placed in an open field arena with one odor in each of the quarters. In addition to the conditioned odors, two neutral odors were presented. The animal was given 10 min twice to explore the arena with an inter-trial time of 10 min in between. Total investigation time was measured in two regions of interest (ROI). First, circular ROIs, corresponding to the odor location, were placed above each petri dish. Second, rectangular ROIs were used, to analyze time spent in each quarter of the arena.

### Head direction-dependent optogenetic stimulation

Animals were put in a cylindrical white arena with a single cue (a black vertical bar), limiting further cue-based orientation by surrounding the arena with black curtain. A random point was chosen to act as a reference for head direction (0°). The reference point was kept constant between experimental sessions and animals but was not visible to the animal. To habituate the animal to the arena, the animal was put into the arena for 30 min for 2 days and reward pellets were placed randomly inside the arena at the 0, 10 and 20 min mark. During the experiment, light stimulation (488 nm, 15 mW) was initiated whenever the animal’s head direction was within a 60° window around the reference point. Stimulation lasted 1 second or as long as the head direction was maintained in the window up to a maximum of 5 seconds. After each stimulation, further stimulation was discontinued for at least 15 seconds to avoid over-heating of brain tissue. The animal was allowed to investigate the arena over 5 consecutive days for 30 min sessions each day during which the animal was stimulated. Animals were perfused 2 days after the last session.

### Action-dependent, light stimulation

The head direction of the animal was measured constantly using the nose and neck point. Whenever the animal was within a 60° window of the reference point, a 488 nm laser (OBIS LX/LS, Coherent Inc., Santa Clara, CA USA) connected to a laser remote (OBIS LX/LS Single Laser Remote, Coherent Inc., Santa Clara, CA USA) was triggered. Each stimulation event (15 mW) lasted at least 1 second or until the animal’s head left the angular window but never longer than 5 seconds. After a delivery of a light stimulus, there was a built in inter-stimulus time of 15 sec.

### Experimental Setup

The corresponding arenas were placed in a closable compartment with isolation from external light sources. A light source was placed next to the setup so that the arena was evenly lit. The camera was placed directly above the arena. During experiments, the compartment was closed to minimize any disrupting influences from outside. All devices were triggered using NI 6341 data-acquisition board (National Instruments Germany GmbH, Munich) in combination with the Python *nidaqxm* library connected via USB 3.0 to a PC (Intel Core i7-9700K @ 3.60GHz, 64 GB DDR4 RAM and NVidia GeForce RTX 2080 Ti(12GB) GPU). For all experiments, we used the Intel Realsense Depth Camera D435 (Intel Corp., Santa Clara, CA, USA) at 848 × 480 and 30 Hz to enable reliable streaming at all times. Although the webcam is capable of 60 Hz and higher resolution, we found that these settings gave reliable framerate and the optional addition of depth data.

We have successfully installed and tested DLStream on Windows 10 and Ubuntu 18.04.05 OS. DLStream was developed in the open-source programming language Python. Python includes open-source libraries for most available devices or desired functions, which allows DLStream to utilize and control a wide range of devices. Virtually any webcam/camera can be used with any framerate and resolution considering hardware requirements and limitations.

### Reward delivery and acoustic stimulation

Liquid reward was delivered via a custom-built reward delivery system using a peristaltic pump (Takasago Electric, Inc.). A nozzle connected to the pump was placed in the corner of the arena. The animal was briefly habituated to the reward and the reward delivery system. The liquid reward consisted of diluted sweetened condensed milk (1:10 with Aqua dest.) and was delivered in a volume of ca. 4-6 µl. If not collected, the reward was withdrawn again. The aversive tone (ca. 100 dB) was delivered via a custom build piezo alarm tone generator. The device was placed above the arena.

### Pose estimation using DLC

In all experiments, we used 3-point tracking to estimate the position, direction, and angle of the animal using the head, neck and tail root as body parts of interest. Networks were trained using the DLC 1.11 framework. First, 300 images of relevant behavior in the corresponding arena were annotated and fed into the DLC network training sets. Second, networks were trained for 500,000 iterations and evaluated. For each experiment type a different network was trained.

### Posture detection in DLStream

We extracted the raw score maps from the deep neuronal network analysis and used them for posture detection. First, body part estimation, similar to the DLC approach, was conducted by local maxima detection using custom image analysis scripts. Resulting pose estimation was then transferred into postures. For this, each possible combination of body parts was investigated and filtered using a closest distance approach. DLStream detects estimated postures and compares them to relevant trigger modules for closed-loop control of experiments.

